# AMPK regulates germline stem cell quiescence and integrity through an endogenous small RNA pathway

**DOI:** 10.1101/455253

**Authors:** Pratik Kadekar, Richard Roy

## Abstract

*C. elegans* larvae can undergo a global developmental arrest following the execution of a diapause-like state called ‘dauer’ in response to unfavourable growth conditions. Survival in this stage surpasses the normal lifespan of reproductive animals quite dramatically, and without any apparent negative impact on their reproductive fitness. During this period, the germ cells become quiescent and must retain their reproductive integrity so the animal can reproduce following recovery. This germline stem cell (GSC) arrest requires the activity of AMP-activated protein kinase (AMPK) and in its absence the germ line undergoes hyperplasia. We show here that AMPK mutant animals exhibit complete sterility after recovery from dauer, suggesting that germ cell integrity is compromised during this stage in the absence of AMPK. These defects correlate with altered abundance and distribution of a number of chromatin modifications that affect gene expression. These aberrant chromatin modifications, along with the supernumerary germ cell divisions and the observed post-dauer sterility, were all corrected by disabling key effectors of the small interfering RNA pathway (*dcr-1* and *rde-4*) and the primary Argonaute protein *ergo-1,* suggesting that AMPK regulates the function of these small RNA pathway components, and in its absence, the pathways become abnormally active. The aberrant regulation of the small RNA pathway components releases the germ cells from quiescence to proliferative state thereby compromising germ cell integrity. Curiously, AMPK expression in either the neurons or the excretory system is sufficient to restore the GSC quiescence and the fertility in the AMPK mutant post-dauer adults, while the fertility of these animals is also partially restored by disabling the dsRNA importer SID-1. Our data suggest that AMPK regulates a small RNA pathway in the soma to establish and/or maintain GSC quiescence and integrity cell non-autonomously in response to the energy stress associated with the dauer stage. Our findings therefore provide a unique model to better understand how the soma communicates with the germ line to establish the appropriate epigenetic modifications required to adapt to acute environmental challenges.

## Introduction

It is becoming more widely accepted that life history can affect developmental and behavioural outcomes, either in a temporary, or often in a more permanent manner. These modifications can occur downstream of a broad spectrum of environmental factors, including temperature, light, resource availability, population density, and even the presence of predators; all of which can influence gene expression, often with dramatic phenotypic consequences (1, 2). Furthermore, these consequences are not restricted to the generation that experienced the event, but rather, they can be transmitted to subsequent generations.

Studies have shown that the molecular record of these events is encoded in the form of epigenetic changes associated with histone modifications, DNA methylation and/or base modification, or alterations in the small RNA repertoire (3). Because the transmission of these molecular memories can span one or several generations, these modifications must impinge in some way upon the germ line, thus providing some adaptive phenotypic change in the unexposed future generations (4-7). These epigenetic modifications in the germ cells can have a significant impact on successive generations, yet the molecular mechanisms through which “experience” is transduced to the genome across several generations remains ill-defined.

*C. elegans* has been used successfully to demonstrate how environmental cues can modulate epigenetic change and behaviour (8). Furthermore, a subset of these modifications and associated traits can be transmitted to subsequent generations in a manner dependent on small heritable RNAs (9). Recently, it was shown that acute starvation at the L1 larval stage leads to the generation of small RNA species that are inherited for at least three generations. This heritable pool of RNAs could reflect the adaptive change in expression of genes involved in nutrition and metabolism (6, 10).

In addition to the L1 stage, later in development, larvae can execute an alternative developmental program to enhance survival and fitness in response to overcrowding or sub-optimal survival conditions. During this diapause-like state called dauer, they undergo a global, genome-wide, adjustment of chromatin modifications that is accompanied by a significant change in gene expression when compared to the animals that never transited through this stage. These changes in the abundance and distribution of chromatin marks likely contribute to the molecular record of life history and the adaptive adjustment of these chromatin modifications is most probably dependent on the expression of specific endogenous small RNAs (11, 12). Currently, it is still unclear how the physiological stress associated with the dauer stage might impact the population of small RNAs and, in transgenerational contexts, how these changes are transmitted across generations, in light of the erasure of histone marks that normally takes place during each cycle of embryogenesis.

Global cell cycle arrest is one of several distinctive features of *C. elegans* dauer larvae. Upon entry into the dauer stage the germline stem cell (GSC) divisions begin to slow to finally establish a state of quiescence, which they maintain until they recover from dauer and resume normal development. Despite potentially long periods in this diapause stage, this cell cycle/developmental quiescence has no impact on their reproductive fitness (11). The activity of the cellular energy sensor AMPK and its upstream kinase LKB1/PAR-4, as well as the activity of tumour suppressor PTEN, are all independently required for the quiescent state of the GSC in response to dauer signalling (13). Maintaining a quiescent state in response to energetic stress may be favourable for survival, presumably because it reduces energy consumption during a period when energy is limited (14, 15). Moreover, when quiescence fails to establish and/or maintained during periods of energy stress, germ cell integrity is compromised, resulting in reduced brood sizes (4).

We show here that during the dauer stage, AMPK activity is not only required to block GSC proliferation, but also to maintain GSC integrity to ensure reproductive success following recovery to replete conditions. In the absence of AMPK, several chromatin modifications become misregulated, resulting in inappropriate gene expression that has a detrimental effect on reproductive fitness following exit from the dauer stage. Using genetic analysis, we reveal the importance of the endogenous small RNA pathway and its regulation by AMPK. Moreover, this pathway acts at least partially in a cell non-autonomous manner, to adjust the GSC-specific chromatin landscape in favour of an adaptive gene expression program fine-tuned toward maintaining germ cell integrity during the long-term energy stress typical of the dauer stage.

## Results

### Defects in the dauer germ line result in post-dauer sterility in AMPK mutants

In *C. elegans,* the decision to execute dauer development is regulated by three independent signalling pathways that converge on a nuclear hormone receptor to ultimately affect multiple developmental and physiological processes (16). Many of these processes involve measures to conserve energy for the duration of the diapause, which are mediated through a significant metabolic adjustment that occurs downstream of all these signalling pathways (17).

To conserve the energy, while also ensuring that germ cell cells do not replicate during this period when key cellular building blocks may be limiting, the *C. elegans* orthologues of LKB1 (*par-4*) and the regulatory and catalytic components of AMPK cooperate to establish cell cycle and developmental quiescence in the germline stem cells (GSC). Animals that have reduced, or lack all AMPK activity (*aak(0)*), undergo pronounced germline hyperplasia due to supernumerary cell divisions that occur prior to dauer entry (13). It is unclear, however, if these extra cells retain their germ cell integrity and are competent to yield functional gametes.

The dauer larva is remarkable in that it can remain in a quiescent state for months longer than it would normally survive while in its reproductive mode. Nevertheless, it can exit this quiescence upon improvement in growth conditions, to resume reproductive development with no compromise of their reproductive fitness, regardless of the duration of the developmental arrest *per se* (18). The germ cells must therefore retain the appropriate information to maintain their totipotency over lengthy periods so that upon recovery from the diapause the animal can still reproduce without any loss in fitness. Since AMPK and LKB1 are critical to block germ cell divisions during the dauer stage we questioned whether the supernumerary germ cells that are produced in *aak(0)* mutants are indeed competent to generate functional gametes and/or embryos. We therefore quantified the brood size of *daf-2* (control) and *daf-2; aak(0)* animals after allowing both of these mutants to recover after remaining at least 24h in the dauer stage. In contrast to control *daf-2* animals that recover from dauer with very little to no negative reproductive consequence, AMPK mutant animals that transit through dauer for 24 hours or more exhibit highly penetrant post-dauer (PD) sterility upon recovery (Fig. 1A, Fig. S2). The brood size of *daf-2* PD adults was not significantly different from *daf-2* animals that never transit through dauer suggesting that passage through the dauer stage has no impact on reproductive fitness provided that AMPK signalling is active.

**Figure 1.**
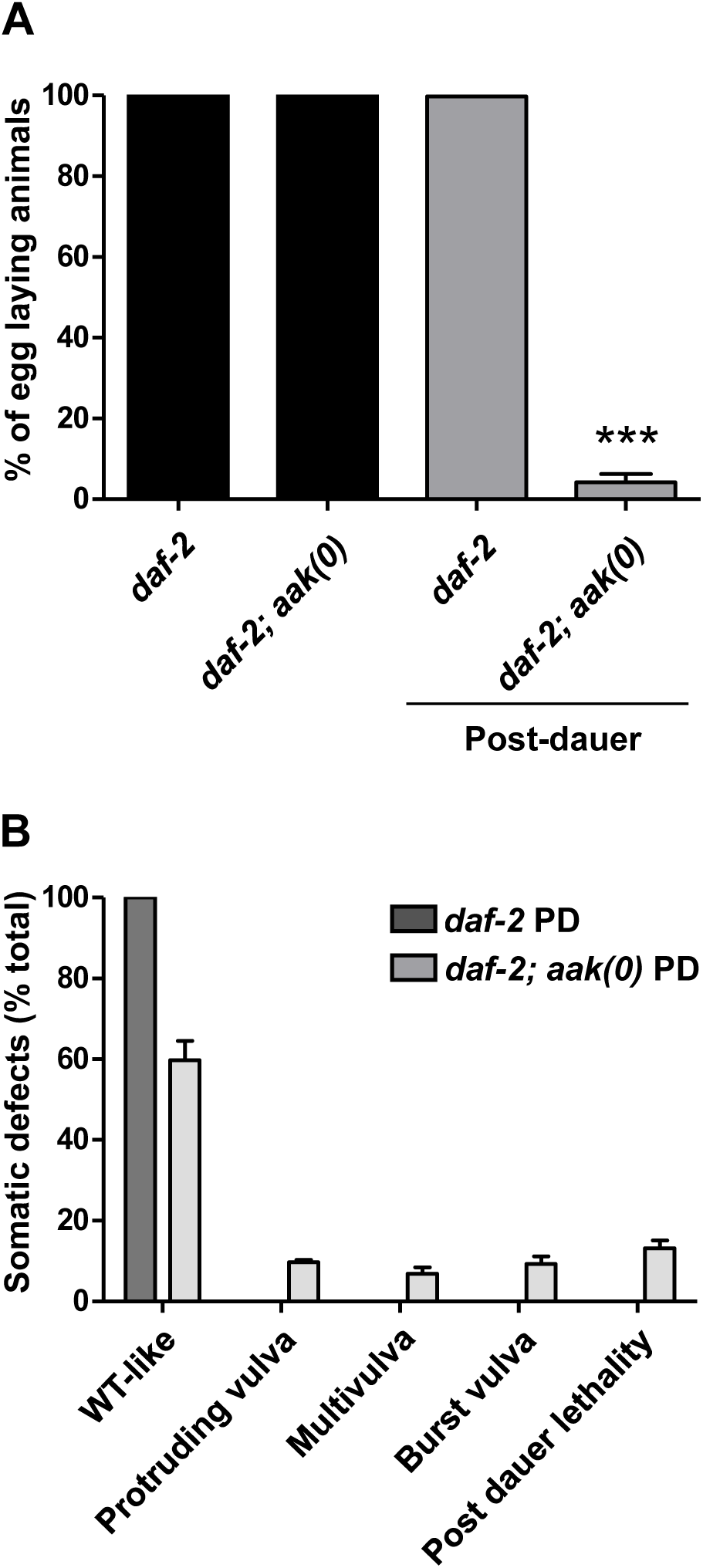
Post-dauer *aak(0)* adults exhibit vulval defects and highly penetrant sterility. **A)** All adult animals that laid eggs were considered as fertile. Both *daf-2* and *daf-2; aak(0)* animals cultivated under permissive conditions showed no fertility defects compared to wild type. To assess the fertility of the PD adults, animals were maintained in the dauer stage for 24 hours after which they were switched to permissive temperature to resume reproductive development (See materials and methods). Egg laying animals were counted, the means calculated, and the values are shown with SD. Upon recovery, *daf-2* PD adults were fertile, but *daf-2; aak(0)* PD adults were almost entirely sterile; ***P<0.0001 using Marascuilo procedure. Assays were performed three times and the data represent the mean ±SD; n=50. **B)** In *daf-2; aak(0)* PD animals, the highly penetrant sterility is also associated with vulval defects. A proportion of these animals (16.5±3.5%) prematurely expired during their recovery phase and failed to reach adulthood. Values represent means ±SD; n=50.

To confirm that the AMPK activity is not exclusive to the insulin-like signalling branch involved in dauer formation, but rather, it is required downstream of the other signalling pathways that control dauer formation, we determined if AMPK may play a more general role in PD fertility by testing if it is also required in *daf-7* mutants (TGF-β pathway), or in the *aak(0)* mutants treated with dauer pheromone (16). Similar to what occurs in the insulin-like signalling mutants, inducing the dauer stage through the compromise of the TGF-β pathway, or treatment with dauer pheromone, also results in highly penetrant sterility in PD animals that lack AMPK (Fig. S1). This suggests that the activity of AMPK is critical for PD fertility, and hence the maintenance of germ cell integrity, downstream of the major pathways required for dauer formation.

Although most of the *aak(0)* PD animals become sterile, a significant portion of die. Those that survive also show diverse abnormalities in vulva development (Fig. 1B, Table S1). Thus, it is not only the germ line which is compromised in PD animals that lack AMPK, but at least some somatic tissues are also sensitive to loss of AMPK function.

### AMPK post-dauer gonads are morphologically abnormal and the germ cells fail to exit pachytene

To determine the physiological basis of the observed sterility in the *aak(0)* PD animals, we examined their germline morphology and organization using a germ cell membrane marker (19). We noted that in 95% of the animals, the oocyte morphology appeared abnormal and they also lacked the typical, single-file organization seen in the control *daf-2* PD animals (Fig. 2A, B, Table S2). Also, in 60% of the *aak(0)* PD animals, the general gonad symmetry was abnormal in terms of size and shape of the gonadal arms. (Table S2).

**Figure 2.**
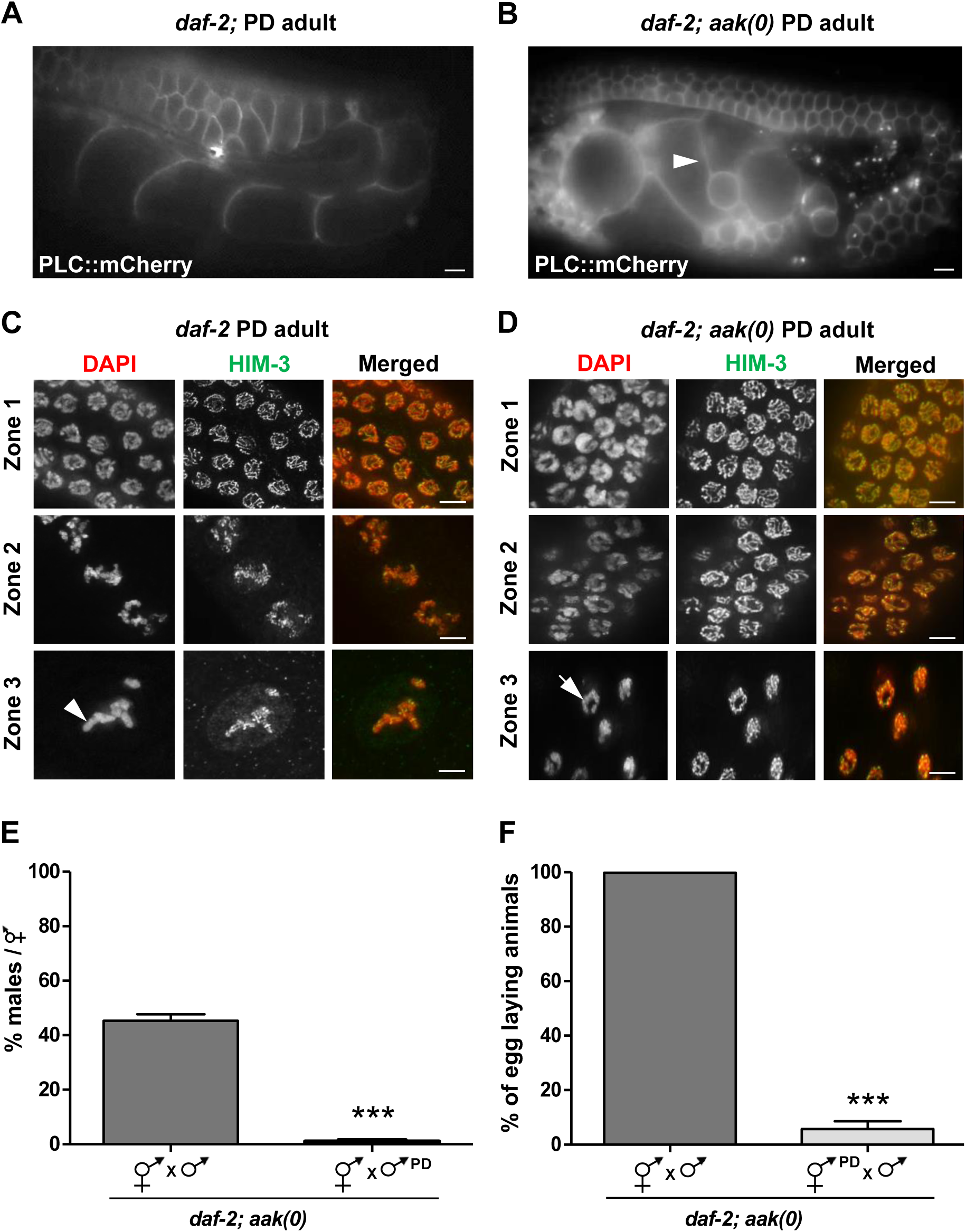
AMPK-defective post-dauer adults show abnormal gonadal morphology and the germ cells fail to exit pachytene. All animals analyzed express a *Ppie-1::* PLC::mCherry transgene to monitor germ cell membranes/organization. **A, B**) In *daf-2* PD adults, the gonad and germ cells develop normally and no obvious defects were observed, but *daf-2; aak(0)* PD adults exhibit various defects in gonad development and organization. Oocyte morphology is abnormal (white arrowhead) and they lacked the typical, file-like organization observed in the control *daf-2* PD animals. **C, D)** For further characterization, the post-transition zone germ cells were divided into 3 different subregions. In the first subregion after the transition zone (Zone 1), germ cells enter pachytene stage; in Zone 2, the cells exit pachytene and initiate the separation of the paired chromosomes (diplotene); in Zone 3 separation of the paired chromosomes is complete, forming 6 tightly condensed DAPI-stained bodies representing 6 pairs of homologous chromosomes (diakinesis). In *daf-*2 PD, the germ cells go through all these processes to eventually give rise to 6 condensed DAPI-stained bodies (white arrowhead), but in *daf-2; aak(0)* PD adults, the germ cells enter pachtytene in Zone 1, but fail to completely exit the pachytene stage based on the continued presence of long chromosome tracks (white arrow). A-D) n= 20. Scale bar: 10um in A and B, 4 um in C and D. **E, F)** Reciprocal crosses were performed and a ratio of 20 males per hermaphrodite was maintained for all the crosses. *daf-2; aak(0)* PD males were mated with normal *daf-2; aak(0)* 15 hermaphrodites and a number of males/hermaphrodite in F1 were counted. Few to no male progeny were identified in the F1 generation of this mating. The mean is shown ± SD. **F)** *daf-2; aak(0)* PD hermaphrodites were crossed with normal *daf-2; aak(0)* males. 15 animals were quantified and PD *aak(0)* hermaphrodites exhibited a high frequency of sterility. The mean is represented ± SD. ***P<0.0001 using Marascuilo procedure.

We stained the germ lines of PD control and AMPK mutant animals and examined chromosome morphology within the germ cells as they progress through the distinct regions of meiotic prophase to generate fully differentiated oocytes (20). There were no obvious defects in the size of the mitotic zone or the spatio-temporal arrangement of the transition zone. For further characterization we binned the post-transition zone germ cells into 3 different zones: in Zone 1, germ cells enter the pachytene stage; in Zone 2 the cells exit pachytene and initiate the separation of the paired homologous chromosomes (diplotene); in Zone 3 separation of homologues is complete, forming 6 tightly condensed DAPI stained bodies representing 6 pairs of homologous chromosomes (diakinesis).

In the *daf-2* PD germline, there were no observed abnormalities as the germ cells complete meiotic progression to eventually give rise to oocytes with 6 condensed DAPI-stained bodies (Fig. 2B). However, in the PD germ lines of *aak(0)* mutants, the germ cells do enter pachytene in Zone 1, but then fail to exit Zone 2. The pachytene arrest persists into Zone 3, as the chromosomes fail to separate and condense (Fig. 2C). This suggests that in *aak(0)* PD animals, the germ cells fail to exit pachytene and thus never undergo diakinesis to produce mature oocytes.

To further determine, if the sterility was a result of abnormal sperm formation or function, or whether the defect was associated with the oocytes, we performed reciprocal crosses and monitored the brood size of the resulting cross progeny (Fig. 2E, F). Using an antibody that detects sperm-specific proteins (anti-MSP), we noted that while sperm was present in the *aak(0)* PD hermaphrodites, it may have been produced during dauer stage (13). To determine if the sperm present in the *aak(0)* PD adults is functional, we mated *aak(0)* hermaphrodites that never transited through dauer, with *aak(0)* PD males (a ratio of 20 males per hermaphrodite was maintained). If the mating was successful and the PD males produced functional sperm, we would expect ~50% of the progeny to be male. However, no significant F1 male progeny were observed suggesting that the sperm is defective (Fig. 2E). We cannot rule out however that, despite our monitoring, these mutants could be mating incompetent.

Similarly, when we mated PD *aak(0)* hermaphrodites and *aak(0)* males that never transited through dauer, *aak(0)* PD hermaphrodites still exhibited highly penetrant sterility (Fig. 2F) suggesting that integrity of the oocytes is also compromised.

These results collectively suggest that germline development is sensitive to periods of energetic stress, and in the absence of AMPK, the germ line becomes severely perturbed; the integrity of both the oocytes and the sperm is affected, ultimately rendering the PD animals sterile.

### Reducing germline hyperplasia does not restore fertility in post-dauer AMPK mutants

As *aak(0)* dauer larvae fail to maintain the GSC arrest and exhibit sterility upon exiting the dauer stage, we questioned whether the observed sterility might result from the inappropriate germ cell divisions that occur during the dauer stage. To test this, we performed RNAi on genes that were shown to suppress the supernumerary germ cell divisions in the *aak(0)* dauer larvae (21). We subsequently allowed these animals to recover to form adults, after which we assessed their fertility and brood size. None of the suppressors of dauer germline hyperplasia that we tested were capable of restoring the PD fertility in the *aak(0)* mutants, although the germline hyperplasia typical of *aak(0)* mutant dauer larvae was visibly ameliorated (Fig. 3A, B). Albeit, RNAi of these genes does not completely suppress the germline hyperplasia, leaving the possibility that the extra number of germ cells could be responsible for the PD sterility. Nevertheless, our data suggest that the PD sterility of *aak(0)* animals is not necessarily a direct consequence of the dauer germline hyperplasia, but may involve additional processes that could act in concert with, but possibly independent of, the regulation of cell cycle quiescence during the dauer stage.

**Figure 3.**
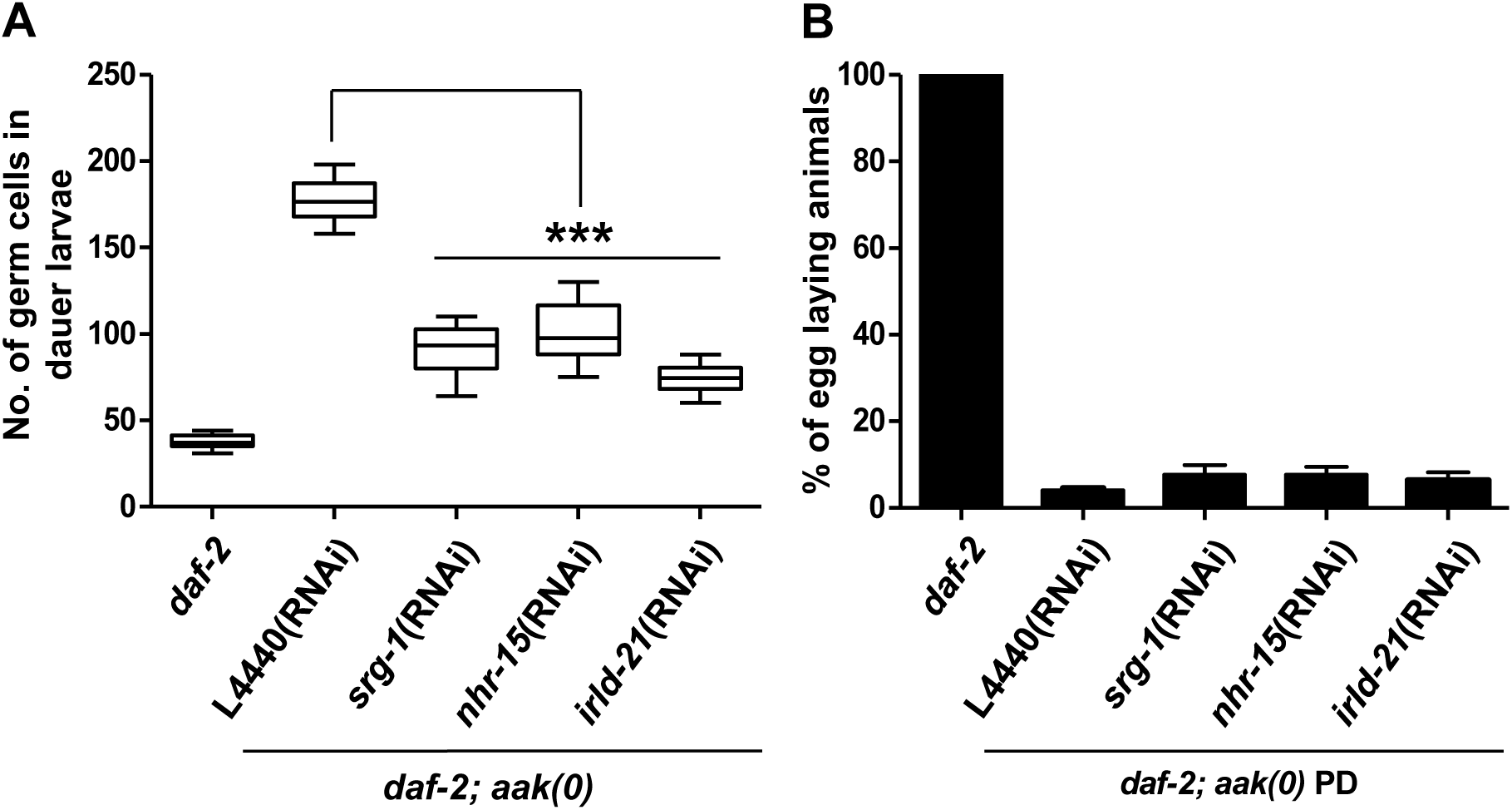
Post-dauer sterility and germline hyperplasia are not obligately interdependent in AMPK mutants. **A, B**) Whole animal DAPI staining was performed to quantify the number of dauer germ cells. The number of egg-laying animals was quantified and the mean is represented ± SD. To test if the germ cell integrity defect results from the dauer-dependent germline hyperplasia, we used RNAi to disrupt three gene products previously found to suppress germline hyperplasia in dauer larvae (21). Larvae were switched to permissive temperature to exit dauer and resume reproductive development. Fertility was assessed 48h after the temperature shift by counting egg-laying adults. L4440 is an empty RNAi vector and is used as a control. ***P<0.0001 when compared with L4440 using the two-tailed t-test. n=50.

### Many chromatin marks are misregulated both globally and in the germ line in *aak(0)* dauer larvae

*C. elegans* dauer larvae exhibit a significantly different gene expression profile when compared to animals that never transit through the dauer stage (11). Furthermore, these changes in the gene expression persist after the animals exit dauer and become reproductive adults. Thus, a molecular memory of the passage through dauer is recorded, and has been shown to influence fertility in the PD animals (11). The observed changes in gene expression are highly correlated with the changes that occur in the various chromatin marks detected in dauer and in PD larvae compared to controls that never transited through dauer (11).

AMPK has been implicated in the regulation of gene expression through its ability to modify chromatin by directly phosphorylating histone H2B to activate stress-promoted transcription (22). Furthermore, recently we showed that AMPK modulates the chromatin landscape to ensure that transcriptional activity is blocked in the primordial germ cells until animals have sufficient cellular energy levels (4). Since AMPK may directly regulate histone modifying enzymes to bring about changes in gene expression we wondered whether chromatin modification may be perturbed in the dauer germ cells, thereby affecting the adaptive gene expression program that would normally occur in dauer. This inability to appropriately adjust to the energy stress associated with dauer development might explain the loss of integrity in the PD germ cells.

We therefore examined the global levels of diverse chromatin marks that were previously found to change following transit through the dauer program. We first performed western blot analysis on whole extracts from *daf-2* and *daf-2; aak(0)* mutant dauer larvae using antibodies specific for histone modifications that are associated both with transcription activation (H3K4me3 and H3K9ac) and repression (H3K9me3 and H3K27me3). Interestingly, all the marks we tested were abnormally high in the absence of AMPK (Fig. 4A). To confirm if the increased level of these chromatin marks was associated with the hyperplasia associated with AMPK mutant dauer larvae, we performed the same experiments in animals that lack *glp-1* (Fig. S3) (20) and quantified the levels of the chromatin marks. The reduction of germ cells significantly decreased the levels of the chromatin marks, suggesting that the germ cells are the major contributors to the global increase in the levels of the chromatin marks in AMPK mutant dauer larvae, although the levels also increase in the soma (Fig. 4B).

**Figure 4.**
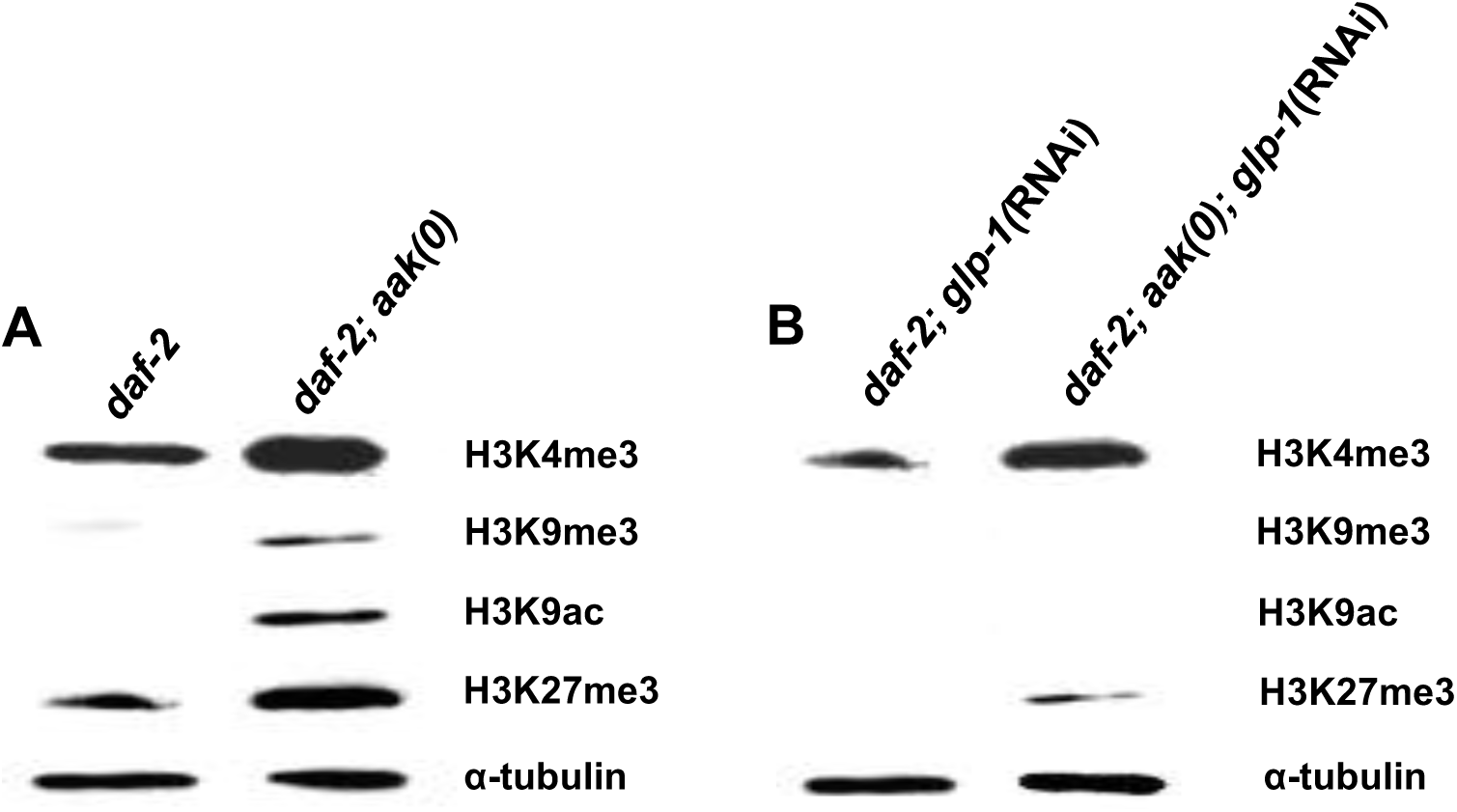
AMPK modulates the abundance of diverse chromatin marks in the soma and the germ line during the dauer stage. **A, B)** Global levels of H3K4me3, H3K9me3, H3K9ac, and H3K27me3 were quantified by performing whole animal western blot analysis of *daf-2* and *daf-2; aak(0)* dauer larvae. *glp-1*(RNAi) was performed post-embryonically using dsRNA feeding in order to compromise germline development without affecting early embryogenesis. α-tubulin is used as a loading control.

### The distribution of chromatin marks is dramatically altered in *aak(0)* dauer germ cells and higher levels of these marks persist in the post-dauer germ line

From our western analysis we could not discern if the levels in the chromatin marks were abundant simply because of the supernumerary germ cells in the AMPK mutant dauer larvae or whether the levels were higher in each individual nucleus *per se.* We therefore dissected gonads from both *daf-2* and *daf-2; aak(0)* dauer larvae and quantified the levels of histone H3 lysine 4 trimethylation (H3K4me3) and histone H3 lysine 9 trimethylation (H3K9me3) to better evaluate their levels per nucleus and to determine if there were any changes in their proximal-distal distribution through the gonad. In the control *daf-2* dauer germ line, the levels of both H3K4me3 and H3K9me3 are uniform across all nuclei throughout the dauer germ line, but in the AMPK mutant dauer larvae, the pattern of H3K4me3 and H3K9me3 expression in the gonad is altered, while their levels are highly variable in individual nucleus (Fig. 5A, B). Of particular interest, we noted that the expression of H3K4me3 in individual nucleus is comparatively weak in the distal gonad, but gradually increases toward the proximal goal where it is much higher.

**Figure 5.**
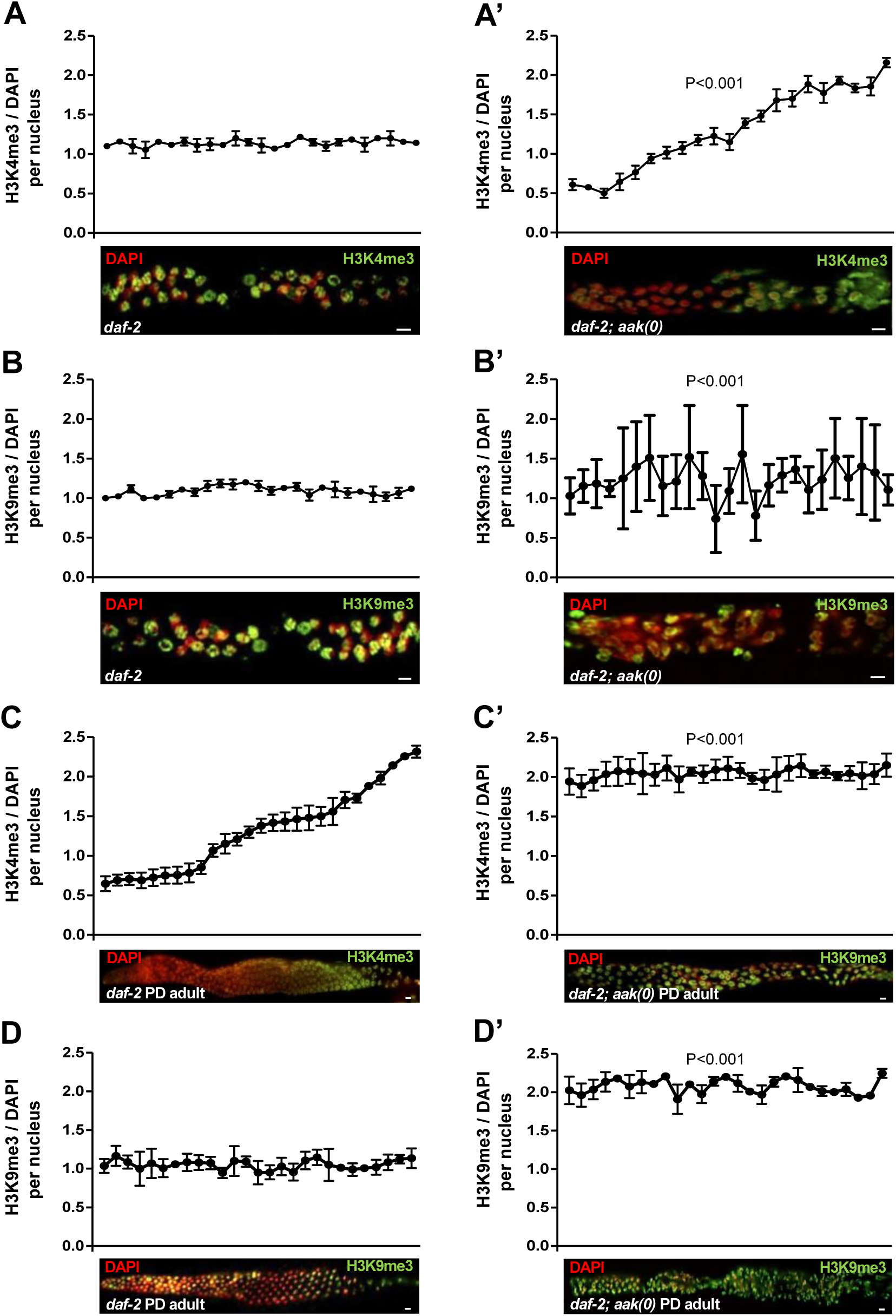
The distribution and abundance of both activating and repressive chromatin marks are dramatically altered in the *aak(0)* dauer and post-dauer germ cells. All images are merged, condensed Z stacks. The graphs represent the average immunofluorescence signal of anti-H3K4me3 and anti-H3K9me3 normalized to DAPI across the dissected germ line. For the micrographs of *daf-2* dauer gonads, the entire dauer gonad was analyzed (distal, proximal, distal). Due to technical difficulties only a single gonadal arm of the *daf-2; aak(0)* gonad was analyzed (distal, proximal). Images in A’, B’, C, C’, D and D’ are aligned such that distal is left side and the proximal is right. **A-A’)** The left panel (*daf-2*) and right panel (*daf-2; aak(0)*) show H3K4me3 (green), and in **B, B’)** H3K9me3 (green) staining merged with DAPI (red). **C-C’, D-D’)** PD *daf-2* and *daf-2; aak(0)* adult gonads were extruded and stained with anti-H3K4me3 and H3K9me3 (green) and signal intensity was quantified throughout the gonad using Image J software. **P<0.001 using the F-test for variance when compared to *daf-2; aak(0)*. Scale bar: 4um n=15 for all the experiments.

To test whether the abnormal distribution and abundance of the H3K4me3 and H3K9me3 marks are resolved after the *aak(0)* larvae exit dauer, we examined these marks in PD adult gonads. Interestingly, we noted that higher levels of both H3K4me3 and H3K9me3 persist in the *aak(0)* PD adult germ line when compared to the control *daf-2* PD animals (Fig. 5C, D).

Altogether, these data confirm the role of AMPK in the regulation of both transcriptionally activating and repressive chromatin marks in the germ line under energetic stress. In its absence the levels of each mark we tested increased, while the distribution of these marks was dramatically disrupted.

### Gene expression is altered in both the *aak(0)* dauer and post-dauer germ line

To examine if aberrant chromatin modifications result in abnormal gene expression in animals that lack AMPK signalling, we performed qRT-PCR to quantify the transcript levels of selected germline-specific genes which were found to be significantly altered in dauer and in the PD adults compared to animals that never transited through dauer (11). The abundance of these transcripts was considerably different in *aak(0)* dauer larvae; some of the genes (*prk-2, pmk-1, mek-2)* were present at lower levels, while others (*spe-26, pro-2)* were detected at significantly higher levels when compared to control *daf-2* animals (Fig. 6A). Furthermore, some of these differences in transcript abundances are not resolved in the PD adult germ line of AMPK mutants (Fig. 6B). Therefore, the chromatin modifications that we observed in both the dauer and PD animals that lack all AMPK signalling result in dramatic deviation from the gene expression program that would normally occur in the germ line as a result of transit through the dauer stage.

**Figure 6.**
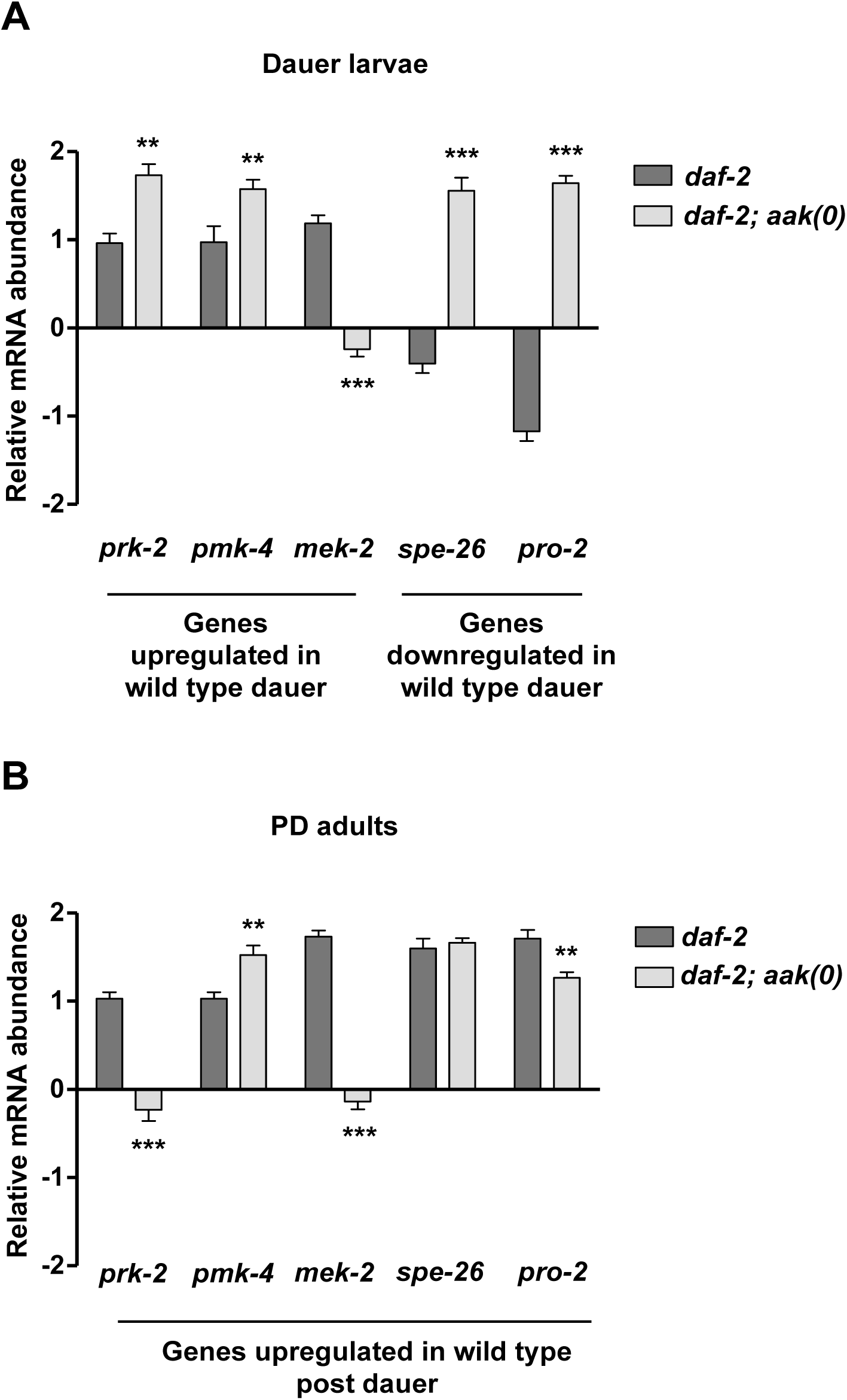
Gene expression is altered in both the *aak(0)* dauer and post-dauer germ line. **A, B)** Germline genes which were previously shown to be differentially expressed during and after transit through the dauer stage (Hall, Beverly et al. 2010), were quantified in *daf-2* and *daf-2; aak(0)* dauer and PD animals. The relative mRNA levels were analyzed using quantitative real-time PCR in both *daf-2* and *daf-2; aak(0)* dauer and PD adults. The expression of these germline genes was significantly altered in *daf-2; aak(0)* dauer and PD animals, when compared to *daf-2*. Error bars, indicate SD from 3 independent experiments. **P<0.001 using one-way ANOVA when compared to *daf-2*.

### Compromise of endogenous siRNA pathway components partially suppresses *aak(0)* post-dauer sterility and dauer germline hyperplasia

Many endogenous small RNAs are critical in distinguishing loci to be targeted by chromatin modifying enzymes, in addition to specifying whether the modification will be active or repressive. They act as mediators of gene expression in order to adapt to cell type information and to varying environmental situations (23). It is therefore not surprising that the small RNA repertoire is dramatically altered in both dauer and PD adults compared to animals that develop in a replete environment (12). We were therefore curious to know whether AMPK might regulate the changes that occur in the suite of small RNA species during dauer owing to its role in regulating chromatin marks in response to energetic stress. To test this possibility, we compromised critical components of the miRNA (*ain-1*), and germ line/nuclear RNAi (*hrde-1, csr-1*) pathways, and some common upstream effectors that impinge on all the small RNA pathways (*dcr-1, rde-4*), to determine if disabling any of these pathways might affect the sterility of the PD AMPK mutant adults.

The miRNA pathway is essential for executing dauer entry making it difficult to interpret their potential role in this process (24). On the other hand, we found that the individual disruption of the RNAse III-like Dicer (*dcr-1*), its accessory factor RDE-4, or the primary Argonaute protein ERGO, could partially rescue the sterility of PD AMPK mutants, the uncontrolled proliferation in the germ line of AMPK mutant dauer larvae, in addition to some of the somatic defects (Fig. 7A, B, Fig. S2, Table S1). In contrast, the compromise of the nuclear Argonaute proteins HRDE-1, or the germline licensing Argonaute CSR-1, had little effect on the hyperplasia or the PD sterility of the AMPK mutants. These data indicate that AMPK must directly, or indirectly, regulate an endogenous small RNA pathway that affects both GSC proliferation and integrity, but that does not include the canonical nuclear Argonaute proteins that have been characterized to regulate germline gene expression.

**Figure 7.**
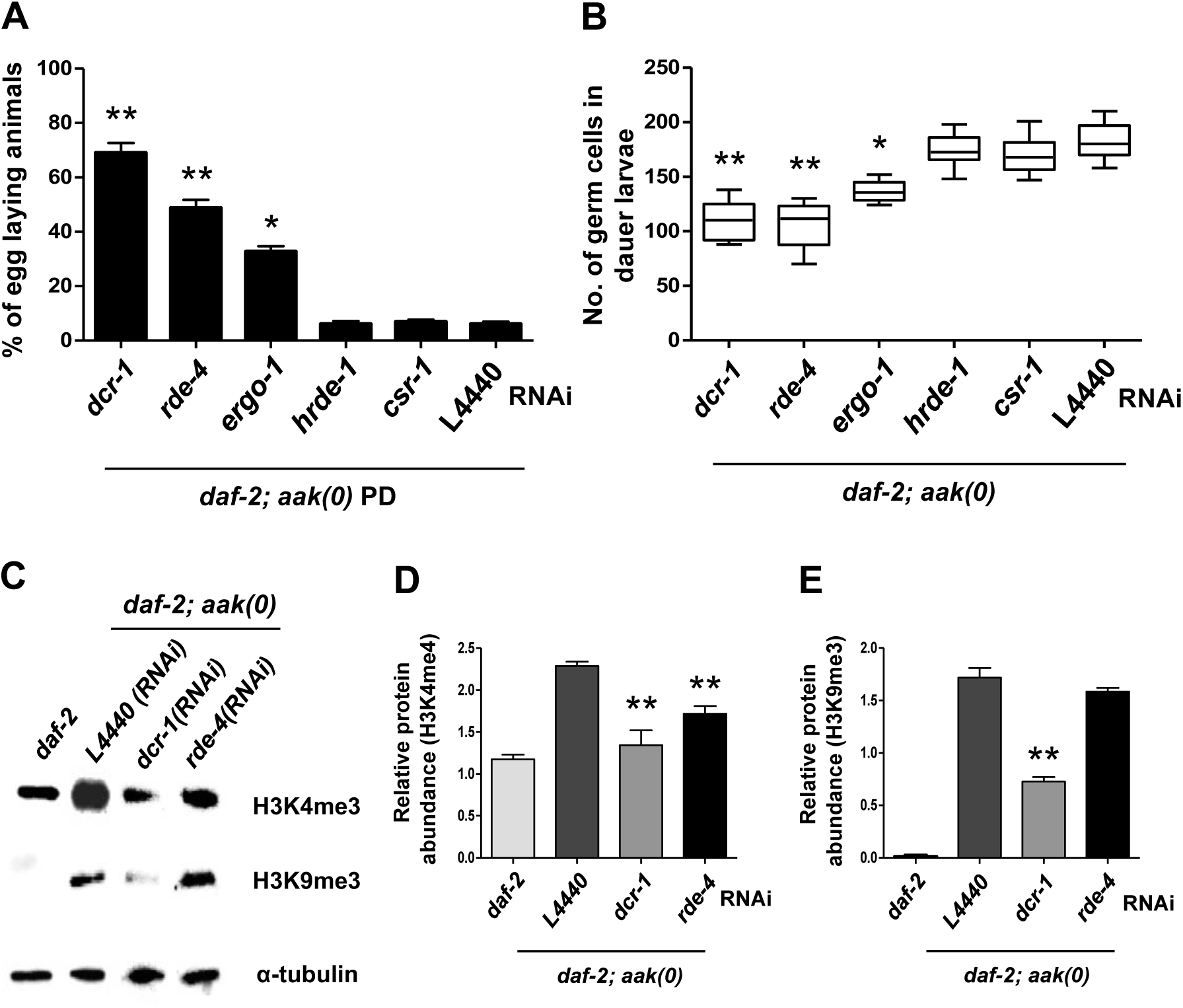
Compromise of small RNA pathway components partially suppresses *aak(0)* post-dauer sterility and dauer germline hyperplasia. To compromise the function of small RNAi pathway, *daf-2; aak(0)* animals were subjected to RNAi by dsRNA feeding against multiple components of the endogenous RNAi pathway. The L4440 empty RNAi vector was used as a control. **A)** The PD sterility observed in the *daf-2; aak(0)* animals was partially rescued by *dcr-1, rde-4* and *ergo-1* RNAi, while RNAi for the germline nuclear Argonautes, *csr-1* and *hrde-1* failed to suppress the observed sterility. **P<0.001 and *P<0.05 using Marascuilo procedure and n=100. **B)** Whole animal DAPI staining was performed to quantify the number of germ cells and the germline hyperplasia in the *daf-2;* aak*(0)* dauer larvae. Statistical analysis was performed using the two-tailed t-test when compared to L4440 treated animals where **P0.001 and *P<0.05; n=100. **C, D, E)** Following the RNAi treatment, global levels of H3K4me3 and H3K9me3 were quantified using whole animal western analysis. Global levels of H3K4me3 were significantly decreased in both *dcr-1* and *rde-4* compromised dauer animals.

Since both the dauer-associated hyperplasia and the PD sterility of the AMPK mutants correlated with the misregulation of chromatin modifications, we wanted to confirm if the compromise of these various RNAi pathway components might also restore the inappropriate levels and distribution of the chromatin marks in the AMPK mutant germ line. We therefore quantified the global levels of both H3K4me3 and H3K9me3 in *dcr-1*(RNAi) and *rde-4*(RNAi) treated dauer larvae. The levels of both of these marks were significantly reduced in the *dcr-1*(RNAi) AMPK mutant dauer animals, with no effect in control *daf-2* dauer larvae. Surprisingly, only the level of H3K4me3 was significantly reduced in the *rde-4*(RNAi) AMPK mutants (Fig. 7C, D, E), although this may reflect the weak and variable RNAi penetrance typical of *rde-4*. These data confirm that AMPK impinges on the endogenous small RNA pathway to modulate chromatin modifications that affect both GSC proliferation and integrity, although it is currently unclear how AMPK might control these processes, and whether phospho-regulation of key targets might be involved.

### Somatic AMPK activity is sufficient to regulate dauer germ cell quiescence and integrity through small RNA transmission to the germ line

Recent studies have shown that AMPK can act cell non-autonomously to regulate lifespan, and the L1 survival of AMPK mutants is greatly improved when AMPK is restored in neurons (25, 26). To determine if AMPK plays a non-autonomous role in maintaining GSC quiescence and integrity, we expressed the catalytic subunit of AMPK (*aak-2*) ubiquitously in the soma (*sur-5p)* of *aak-2* mutants. Interestingly, ubiquitous somatic expression of *aak-2* restores the fertility in the PD of *aak-2* mutants and also rescues the dauer germline hyperplasia (Fig. 8A, B and Fig. S2). This suggests that AMPK activity in the soma is sufficient to maintain the integrity and quiescence in the germ cells during the dauer stage. To determine in which somatic tissue AMPK function can restore quiescence and integrity to the GSCs during the dauer stage, we used tissue-specific promoters to express *aak-2* and quantified PD sterility and the degree of germline hyperplasia in the AMPK mutants. Using a transgenic strain collection, we generated AMPK null mutants that express *aak-2* exclusively in the neurons (*unc-119)*, the excretory system (*sulp-5)*, the skin (*dpy-7*), the gut (*elt-2*), and the muscles (*unc-54*) (27). Surprisingly, only the restoration of AMPK function in the neurons and the excretory system could partially rescue the fertility and GSC quiescence (Fig. 8A, B). These observations suggest that AMPK expression in either (or both) of these two tissues is sufficient to control germ cell homeostasis cell non-autonomously during the energetic stress typical of the dauer stage.

**Figure 8.**
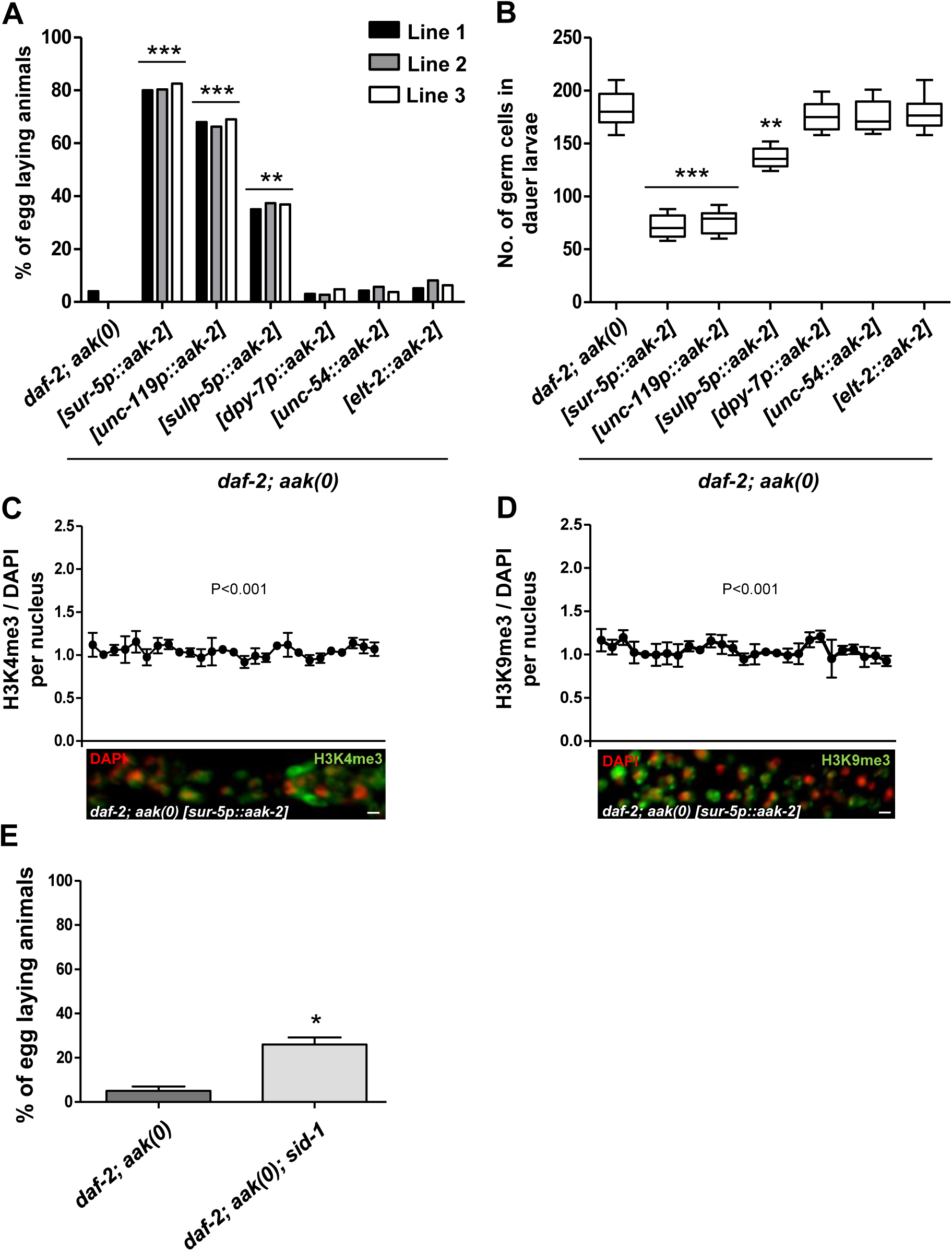
Somatic AMPK activity is sufficient to restore germ cell quiescence and integrity through the transmission of small RNAs in *aak(0)* mutants. **A)** Plasmid constructs that contain *aak-2* cDNA driven by tissue-specific promoters was injected into *daf-2; aak(0)* mutants and both the dauer-dependent germline hyperplasia and the PD sterility were evaluated for each transgenic strain. All transgenic lines are extrachromosomal and are represented by square brackets, and 3 independently generated lines were used for quantification. PD fertility was assessed 24h following the temperature shift after animals were maintained minimally 24 hours in dauer. ***P<0.0001 and **P<0.001 using Marascuilo procedure when compared to *daf-2; aak(0)* and n=50. **B)** Whole animal DAPI staining was performed to quantify the number of germ cells present in the dauer gonad in the transgenic lines and compared to controls. ***P0.0001 and **P<0.001 using the two-tailed t-test when compared to *daf-2; aak(0)* and n=50. **C, D)** All the analyzed images are merged, condensed Z stacks. The graphs represent the average immunofluorescence for H3K4me3 and H3K9me3 normalized to DAPI across the dissected gonad. **P 0.001 using F-test of variance when compared to *daf-2; aak(0)* and n=10. **E)** Disrupting soma to germline transmission of double-stranded RNA by compromising the function of *sid-1* partially restores fertility in the *daf-2; aak(0)* PD animals. A number of animals laying eggs were counted and the mean is shown ± SD. *P<0.05 using Marascuilo procedure when compared to *daf-2; aak(0)* and n=100.

Ubiquitous somatic expression of *aak-2* also restores the normal levels and distribution of H3K4me3 and H3K9me3 marks in the dauer germline (Fig. 8C, D). These data confirm that the somatic function of AMPK can re-establish quiescence and maintain germ cell integrity through its effects on the germline chromatin landscape in dauer larvae that otherwise lack all AMPK signalling.

To further investigate if AMPK modulates some aspect of soma to germ line communication through the deployment of somatically-derived small endogenous RNAs, we tested whether the dsRNA importer, *sid-1,* could affect the PD sterility of AMPK mutant by impairing the uptake of these molecules (28). Interestingly, loss of *sid-1* partially restored fertility in AMPK PD animals (Fig. 8E), suggesting that an AMPK-dependent control switch becomes misregulated in AMPK mutants, allowing the transfer of an aberrant population of small endogenous RNAs to the germ line, culminating in the establishment of an inappropriate chromatin landscape. The resultant gene expression may therefore not reflect the necessary metabolic adjustment that must occur during the dauer stage, altering the ability of the germ cells to adapt to the energy stress associated with dauer. This ultimately compromises their integrity and manifests as the consequent PD sterility.

## Discussion

During periods of energetic stress, *C. elegans* larvae can alter their normal reproductive development and enter a quiescent diapause-like state called dauer. The dauer stage is associated with exceptional stress resistance that is accompanied by a global developmental arrest, including a temporary attenuation of germ cell divisions. One of the predominant cellular energy sensors, AMPK, becomes highly active during this state and regulates the germline quiescence typical of this diapause stage. In the absence of AMPK, or its upstream activating kinase PAR-4/*LKB1,* the dauer germ cells proliferate abnormally, resulting in a dramatic over-proliferation of the germline (13). The consequences of these unscheduled germ cell divisions have never been interrogated. If these cells are competent, the excessive proliferation could result in a significant increase in reproductive fitness. Alternatively, if these supernumerary cells are abnormal it could have detrimental effects on subsequent generations.

We show that the supernumerary germ cells that arise during dauer in AMPK mutants are incompetent to generate functional gametes and therefore provide no reproductive advantage. AMPK signalling is therefore critical to coordinate germ cell quiescence and the appropriate metabolic adjustment required to survive the, often lengthy, organismal energy stress associated with dauer. Although both the dauer germline quiescence and the PD fertility are AMPK dependent, our data suggest that the observed sterility may not necessarily be a direct consequence of abnormal cell divisions within the germline stem cells, since the aberrant cell divisions can be suppressed without ameliorating the sterility of the AMPK mutant dauer larvae. The AMPK-dependent processes that are required for PD fertility may be independent of its role in modulating germ cell proliferation (Fig. 3).

Following dauer recovery, *daf-2* animals develop normally with no consequence on their reproductive fitness. However, AMPK PD animals show striking defects in germline development and organization (Fig. 2B, Table S2). Similar to MAPK signalling mutants, the germ cells fail to completely exit from pachytene-like state in PD animals that lack AMPK (Fig.2) (29). Although, morphologically many of the proximal germ cells appear to undergo cellularization and oogenesis, at the nuclear level they fail to complete diakinesis to form 6 condensed nuclear bodies; a hallmark of a mature oocyte and thus fails to produce a functional matured oocyte (30). AMPK has been shown to regulate the MAPK pathway to block the germ cell divisions under nutritional stress (31). Therefore, perhaps during the energy stress associated with the dauer stage, AMPK could regulate key components of the MAPK pathway to modulate meiotic cell cycle progression thereby delaying the production of mature oocytes, and ensuring that reproductive development is coordinated with transit through the dauer stage.

This interaction with the MAPK pathway might also account for some of the vulval defects that arise in PD mutant animals, although it is not clear what component is affected in either of these contexts. Curiously, like the germ cells, the vulval precursor cells must also maintain an undifferentiated state during the entire length of dauer, only to re-initiate their divisions and fate specification in response to recovery cues (32, 33). Perhaps AMPK is critical for the appropriate maintenance of developmental plasticity in these somatic cells in a manner that is akin to its role in the germ line. Collectively, these results suggest the role of AMPK activity in establishing GSC cell cycle quiescence, meiotic progression, and maintaining germ cell integrity during periods of extreme energetic stress.

But what might constitute germ cell integrity and how might it be maintained by AMPK over the duration of the dauer stage? One possibility might include changes in the gene expression program to mediate the adaptive cellular and metabolic adjustments necessary to endure the energetic stress of the dauer stage, whether it lasts 24h or 6 months. As *C. elegans* larvae enter the dauer stage, the chromatin is concomitantly remodelled, altering gene expression significantly (11, 34). These modifications are tightly correlated with changes in the small RNA repertoire such that the expression of most endo-siRNAs are affected in both dauer and PD larvae when compared with animals that never transit through the dauer stage. Based on the mechanism of small RNA-mediated changes to the chromatin, these changes in small RNA population likely presage chromatin remodelling, which together provide a molecular memory of this life history event, possibly providing a template for the consequent establishment of distinct adaptive cellular responses or behaviour(s) (11).

We show that AMPK is critical to ensure that these global chromatin modifications occur in a regulated manner. In its absence, the abundance of both the transcriptional activating (H3K4me3 and H3K9ac) and repressive (H3K9me3 and H3K27me3) marks increase aberrantly in the soma and the germ line of dauer larvae. The normal distribution of the H3K4me3 and H3K9me3 chromatin marks become visibly perturbed within the AMPK mutant dauer germline. Moreover, the aberrant marks fail to resolve upon dauer exit, persisting in to the adult PD germline, consistent with AMPK further acting to modulate these modifications upon dauer exit. Although we have not identified the penultimate AMPK target(s) that mediate these changes in chromatin regulation, the relationship between AMPK and chromatin regulators may be akin to its role during the L1 diapause, where germ cell integrity is compromised due to irregular chromatin modifications that take place in the primordial germ cells in the absence of AMPK (4).

The observed anomalies in both the abundance and the distribution of the activating and repressive marks likely perturbs the coordination of germline gene expression with the energy stress of dauer, which would normally be mediated through a specific chromatin syntax in both dauer and PD AMPK animals. In the absence of AMPK, gene expression may no longer correspond to that of a germ cell, or at least a germ cell that has been subjected to the challenge of surviving severe energy stress. The inability of the gene expression program to adjust to the metabolic challenge is likely to be responsible for the abnormal gonad and germline development in addition to the somatic defects observed in the AMPK PD adults.

The somatic and germ line abnormalities that we observe in AMPK mutants are dependent on various components of a small RNA pathway. Although small RNA pathways have been previously linked to transcriptional and chromatin modification, our data indicate that these regulatory mechanisms are under direct or indirect control of AMPK during periods of energy stress. Although we have not yet identified the key AMPK targets that mediate this small RNA function, it is noteworthy that Dicer contains multiple consensus AMPK phosphorylation sites, while RDE-4 also could be a potential substrate. Alternatively, in addition to the primary ARGONAUTE protein ERGO, a number of ARGONAUTE orthologues remain to be characterized. We cannot rule out that one of these ARGONAUTE family members may somehow respond to AMPK signalling to affect this small RNA-mediated change in the chromatin landscape that occurs during dauer and PD recovery in *C. elegans*.

Using *rrf-1* to address where AMPK is required, we initially concluded that AMPK acted in a germline-autonomous manner to maintain GSC quiescence in the dauer larvae (13). The technical shortcomings of this strategy have since been well documented (35), and our recent transgenic experiments confirm that AMPK activity is sufficient in the neurons, or the excretory system, to regulate germ cell quiescence and integrity. Moreover, the somatic expression of *aak-2* also restored the chromatin modifications to wild type levels and a normal pattern of distribution throughout the dauer germline. This neuron-specific function of AMPK may not be a general feature of AMPK signalling, since its neuronal expression failed to suppress the supernumerary germ cell divisions in AMPK mutants during L1 diapause (25). Recently, it was shown that AMPK expression in the neurons can extend lifespan in *C. elegans* under energetic stress (25, 26). The neurons may therefore sense the environment and accordingly signal, perhaps in a neuroendocrine manner, to other tissues in order to adapt synchronously as an organism. AMPK could be one of the intermediaries in transducing the signals from the neurons, potentially regulating some diffusible molecule, to enhance their survival without any compromise on their fitness. This would place AMPK at a critical position in sensing environmental challenges to ultimately impinge on the germ line to give rise to the observed chromatin-mediated adaptations associated with dauer, and subsequent recovery in replete conditions.

But what neuron-derived diffusible signal could affect the chromatin in the germ cells? Like many plants, RNAi is systemic in *C. elegans.* Injection of dsRNA into the somatic tissue can result in RNA-mediated gene silencing in the germ line (28, 36, 37). Furthermore, the endo-siRNA pathway that is active in the somatic tissues can contribute, at least in part, to the changes in germline gene expression and the brood size upon passage through dauer (12). Our results support and extend these findings, as the compromise of the dsRNA importer, *sid-1* partially rescues the AMPK-dependent sterility of PD AMPK mutants, suggesting that the abnormal transfer of small RNAs that occurs in the absence of AMPK is registered in the germline culminating in PD sterility. At present our data cannot discern if AMPK regulates the systemic transfer of small RNAs directly, or whether it is involved in selecting the appropriate sequence-specific small RNAs to ensure the correct adjustment to gene expression is achieved during the dauer stage.

The stress associated with the dauer stage is documented molecularly in the chromatin modifications that govern gene expression in PD animals. How this modification in transcriptional output provides some adaptive advantage has not yet been unequivocally determined, but animals that transit through dauer live longer and have a significantly higher brood size (11). These changes may be heritable since animals that remain in dauer for extended periods also show significant alterations in their gene expression, while also demonstrating and enhanced resistance to starvation in subsequent generations (38). Therefore, the dauer larva may be a perfect example of how perceived environmental duress is transduced to the germ line. Most importantly, this may occur through a somatic sensing mechanism that could include AMPK and its ability to modulate diverse stress-specific epigenetic changes via the endogenous small RNA pathway.

At the turn of the last century August Weismann postulated that the germ line was exclusively responsible for the heritable nature of specific traits with little or no impact from the soma. Although this view is widely accepted, diverse situations have been described where the soma acts as a critical regulator of epigenetic change with phenotypic effects that can last for multiple generations (28, 39, 40). AMPK may be one of the critical somatic effectors required to bridge the proposed Weismann Barrier, providing an efficient means of coordinating epigenetic change in the germ line with physiological or environmental cues sensed by the soma. Our findings therefore provide a means of dissecting the mechanisms through which the soma communicates with the germ line in order to adapt to acute environmental challenges, through the generation of a suite of chromatin modifications that confer an epigenetic-based selective advantage to future generations.

## Materials and Methods

### *C. elegans* genetics

All *C. elegans* strains were maintained at 15°C and according to standard protocols (41). The strains used for the study include CB1370 [*daf-2(e1370 III*], MR1000 [*daf-2(e1370) aak-1(tm1944) III; aak-2(ok523) X*], MR0480 [*daf-7(e1372) III; aak-2(ok523) X*], MR1175 [*aak-1(tm1944) III; aak-2(ok523) X*], MR2137 [*daf-2(e1370) aak-1(tm1944) III; aak-2(ok523) X; ltIs4[unc-119(+)Ppie1::plc::mCherry]*], MR2138 [*daf-2; ltIs44[unc-119(+)Ppie1::plc::mCherry]*], MR1973 [*daf-2(e1370) aak-1(tm1944) III; aak-2(ok523) X; sid-1(rr167) V*]. Transgenic lines and compound mutants were created in the laboratory using standard molecular genetic approaches. To create transgenic lines to express tissue-specific *aak-2,* MR1000 animals were injected with different constructs as per (27).

### RNAi Feeding

Bacterial clones expressing dsRNA from the RNAi library were grown in LB medium with ampicillin at 37°C overnight. The bacterial culture was seeded onto regular NGM plates containing ampicillin and IPTG. Seeded plates were incubated at room temperature for 24 hours to induce dsRNA expression. L4 larvae were fed on the RNAi plates and were allowed to lay eggs at 15°C and then the eggs were switched to 25°C to induce dauer formation.

### DAPI staining and counting germ cell nuclei

For whole animal DAPI (4’,6-diamidino-2-phenlindole) staining, dauer larvae were washed off plates and soaked in Carnoy’s solution (60% ethanol, 30% acetic acid, 10% chloroform) on a shaker overnight. Animals were washed twice in PBST (1XPBS + 0.1% Tween 20), and stained in 0.1 mg/ml DAPI solution for 30 minutes. Finally, larvae were washed four times (20 minutes each) in PBST, and mounted in Vectashield medium. The total number of germ cell nuclei per dauer gonad was then determined based on their position and nuclear morphology.

### Dauer recovery assay

A population of the genetically identical animals were synchronized, and the resulting embryos were added to normal NGM plates seeded with *E. coli*, and incubated at 25°C for 72 hours in order to induce dauer formation and allow animals to spend at least 24 hours in dauer state. Following this window, dauer larvae were shifted to the permissive temperature of 15°C to allow dauer larvae to recover and initiate regular development. Upon dauer exit, the L4 larvae were individually isolated onto separate plates and were transferred to new plates every 24 hour intervals to quantify their brood size. The brood size of each animal was the sum of non-hatched and hatched progeny.

### Immunostaining and quantification

For extruded dauer gonad staining, gonads were dissected, fixed and stained as described elsewhere (42). The following primary antibodies were used: rabbit polyclonal anti-H3K4me3 (Abcam, 1:500), anti-H3K9me3 (Cell Signaling Technology, 1:500), rabbit anti-HIM-3 (gift from Zetka lab, 1:200). Secondary antibodies were Alexa Fluor 488-coupled goat anti-rabbit (Life Technologies, 1:500). Microscopy was performed as described in (43). Ratios for the fluorescence intensity across the germ line were determined using Image J.

### Western blot

*C*. *elegans* dauer larvae and PD adults were lysed by sonication in lysis buffer (50mM Hepes pH7.5, 150mM NaCl, 10% glycerol, 1% Triton X-100, 1.5mM MgCl2, 1mM EDTA and protease inhibitors). Protein concentrations were determined using nanodrop 2000c spectrophotometer (Thermo Scientific). Nitrocellulose membranes were incubated with primary antibodies: rabbit anti-H3K4me3 (Abcam, 1:1000), anti-H3K9me3, anti-H3K27me3, anti-H3K9ac (Cell Signaling Technology, 1:1,000); mouse anti–α-tubulin (1:3,000; Sigma). Proteins were visualized using horseradish peroxidase conjugated anti-rabbit or anti-mouse secondary antibody (Bio-Rad).

### RNA Isolation and Real Time PCR

Total RNA was extracted with Trizol (Invitrogen). RNA concentration and purity were determined with a NanoDrop 2000c spectrophotometer. Purified RNA (400 ng) was used to synthesize cDNA. Gene expression levels were determined by real time PCR with the SYBR Green Supermix and BioRad iCycler Real Time PCRSystem (BioRad). Relative gene expression was normalized to *tba-1* which was the loading control.

## Acknowledgements

We are grateful to all Roy laboratory members for their advice and support throughout this work. We acknowledge the CGC for *C. elegans* strains and reagents; Jean-Claude Labbé for sharing the *ltIs44* transgene and Monique Zetka for the HIM-3 antibody. This work was supported by an operating grant to RR from the Canadian Institutes of Health Research (CIHR).

## Author contributions

Experiments were designed by P.K. and R.R. P.K. performed all the experiments and analyses. The manuscript was written by P.K. and edited by R.R.

